# A systematic assessment of single-cell language model configurations

**DOI:** 10.1101/2025.04.02.646825

**Authors:** Gaetan De Waele, Gerben Menschaert, Willem Waegeman

## Abstract

Transformers pre-trained on single-cell transcriptomic data have recently been applied to a series of tasks, earning them the title of foundation models. As all currently published models in this class employ vastly different pre-training strategies, it is impossible to determine which practices drive their success (or failure). Here, we present a large-scale study of pre-training components for single-cell transcriptomic transformers: *bento-sc* (BENchmarking Transformer-Obtained Single-Cell representations). By isolating (and tuning) parts of the pre-training scheme one by one, we define best practices for single-cell language model (scLM) construction. While comparisons with baselines indicate that scLMs do not yet offer the generational leap in prediction performances promised by many foundation models, we identify key design choices leading to their improved performance. Namely, the best scLMs are obtained by: (1) minimally processing counts at the input level, (2) using reconstruction losses that exploit known count distributions, (3) masking (up to high rates), and (4) combining different pre-training tasks/losses. All code supporting this study is distributed on PyPI and is packaged under: https://github.com/gdewael/bento-sc.

## Introduction

Foundation models are large neural networks that are typically pre-trained in a self-supervised fashion (i.e. using a pre-training task that does not require manually-labeled data)^[1]^. Facilitated by the recent construction of large reference cell atlases, they have gained traction as a unifying framework for single-cell transcriptomics (scRNA-seq) analysis^[2;3]^. Often, their promise is to provide *better* performance than purely supervised and specialized counterparts, for a *variety* of tasks, starting from a single model. They can deliver on this promise because their pre-training is supposed to capture intricate patterns in vast unlabeled data^[4]^. Often, they are said to remain useful in “zero-shot” settings, a term which describes the use of the model representations applied to novel tasks without fine-tuning model weights^[5]^.

Transformers arise as a natural architectural candidate for creating foundation models. Their rise to dominance within many subfields of deep learning is partly owed due to their flexible architectural setup^[6]^. The transformer’s self-attention mechanism learns interactions between all input tokens, essentially making them graph neural networks operating on a complete digraph^[7]^. This lack of inductive biases on the patterns they may learn is the reason why they offer supreme representational capabilities when trained on vast data. In addition, biological mechanisms are often represented through graphs, making transformers a conceptually attractive architecture^[8]^. When trained on scRNA-seq data consisting of gene-level counts, transformers learn genegene graphs at every layer. Hence, one may think of self-supervised single-cell transformers as models of the “transcriptomic language”.

Current iterations of single-cell transcriptomic language models primarily consist of transformers pre-trained through masked language modeling (MLM)^[9;10]^. They have been applied to a variety of tasks, including cell-type identification, postperturbation expression prediction, batch correction, and denoising^[11;12;13;14]^. Despite their potential, recent benchmarks indicate that currently available foundation models may often be surpassed by simpler task-specific ones^[15;16;5;17]^.

As all currently published single-cell language models (scLMs) use vastly different model and pretraining designs, it is currently impossible to ascertain which practices drive their success (or failure). Rather than benchmarking the existing models themselves, in this study, we propose to reimplement their design idiosyncracies, and benchmark individual scLM components instead. By doing this, our goal is to outline best practices for scLM construction. To assess language model performance, a comprehensive set of downstream tasks is gathered, covering both (1) cell-as well as gene-level representational tasks, and (2) tasks that involve finetuning or rely purely on inference at test time (zero-shot). Our study design, codenamed *bento-sc* (BENch-marking Transformer-Obtained Single-Cell representations), follows that of a greedy hyperparameter optimization at the scale of large language models. To foster an open space for evaluating future scLM components, all benchmarking and pre-training routines are published as a PyPI-distributed Python package: https://github.com/gdewael/bento-sc.

## Results

### A general recipe for language modeling on single-cell transcriptomics data

Here, we present a general framework for training single-cell language models. This recipe forms the base transformer design upon which all experiments will iterate. In this study, we restrict ourselves to the masked (non-autoregressive) form of language modeling^1^. As a consequence, we use a bidirectional encoder-only transformer. The model counts 51.9M learnable weights. For more architectural details, see Methods.

Following recent work^[11]^, only non-zero counts of the single-cell expression profile are included in the input. Furthermore, all input expression profiles are capped to a maximum of 1024 gene tokens, eliminating the lowest-count genes when this threshold is exceeded. To efficiently train models, the FlashAttention library is used^[19]^. The choice of zero-count exclusion, maximum input genes, and FlashAttention usage is discussed in more detail in Appendix A.

All models are pre-trained using the CELLxGENE census of scTab^[20]^ (scTab dataset). This choice is motivated by its data splits being reproducible, and its history of being used to evaluate self-supervised learning in non-transformer contexts^[4]^. scTab comprises over 20 million human cells covering 19331 human protein-encoding genes.

A general recipe for scRNA-seq MLMs proceeds as follows. Let us denote an expression profile of a single cell by its strictly non-zero (raw) counts: {*c*_1_, *c*_2_, …, *c*_*n*_}, belonging to genes with indices {*i*_1_, *i*_2_, …, *i*_*n*_}, where *n* denotes the number of observed (non-zero) genes in the cell, capped to a maximum of 1024. Gene identity embeddings are obtained through a look-up embedding matrix ***E*** ∈ ℝ^*I*×*d*^, where *I* denotes the total number of unique genes in the dataset (here, 19331) and *d* the embedding dimension. Individual gene ID embeddings are obtained are through 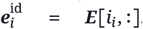. Gene count embeddings are similarly obtained through 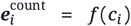, with 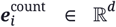 and *f* (·) a count binning and embedding function (see Methods). A final input gene embedding is formed by the sum of identity and count embedding: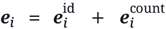. A learnable token vector ***e***_[CLS]_ ∈ ℝ^*d*^ is prepended to the set of gene embeddings for every cell. This token vector serves as a summarization token for the whole cell. A single cell input sample to the transformer model is, hence: ***X*** = [***e***_[CLS]_, ***e***_1_, ***e***_2_, …, ***e***_*n*_], with ***X*** ∈ ℝ^(*n*+1)×*d*^.

During pre-training, a percentage of gene count embeddings 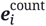 are randomly replaced with a mask embedding ***e***^[MASK]^ at every training iteration. An encoder-only transformer model is then trained to predict what the original pre-masking count values were. It does so by processing the (partly corrupted) input ***X***, and producing (1) a summary cell-level output embedding _***g***[CLS]_, and (2) contextualized output gene embeddings [***g***_1_, ***g***_2_, …, ***g***_*n*_]. Gene embeddings are sent to an expression prediction head (consisting of a position-wise fully-connected layer) returning count predictions ĉ_*i*∈ {1…,*n*}_, which are used to optimize a reconstruction loss w.r.t. the pre-masked counts. The reconstruction loss optimizes the cross-entropy between a predicted probability of a count belonging to a bin and its true count bin. A visual overview of the model is presented in Figure 1. For model and pretraining details, see Methods.

**Figure 1.**
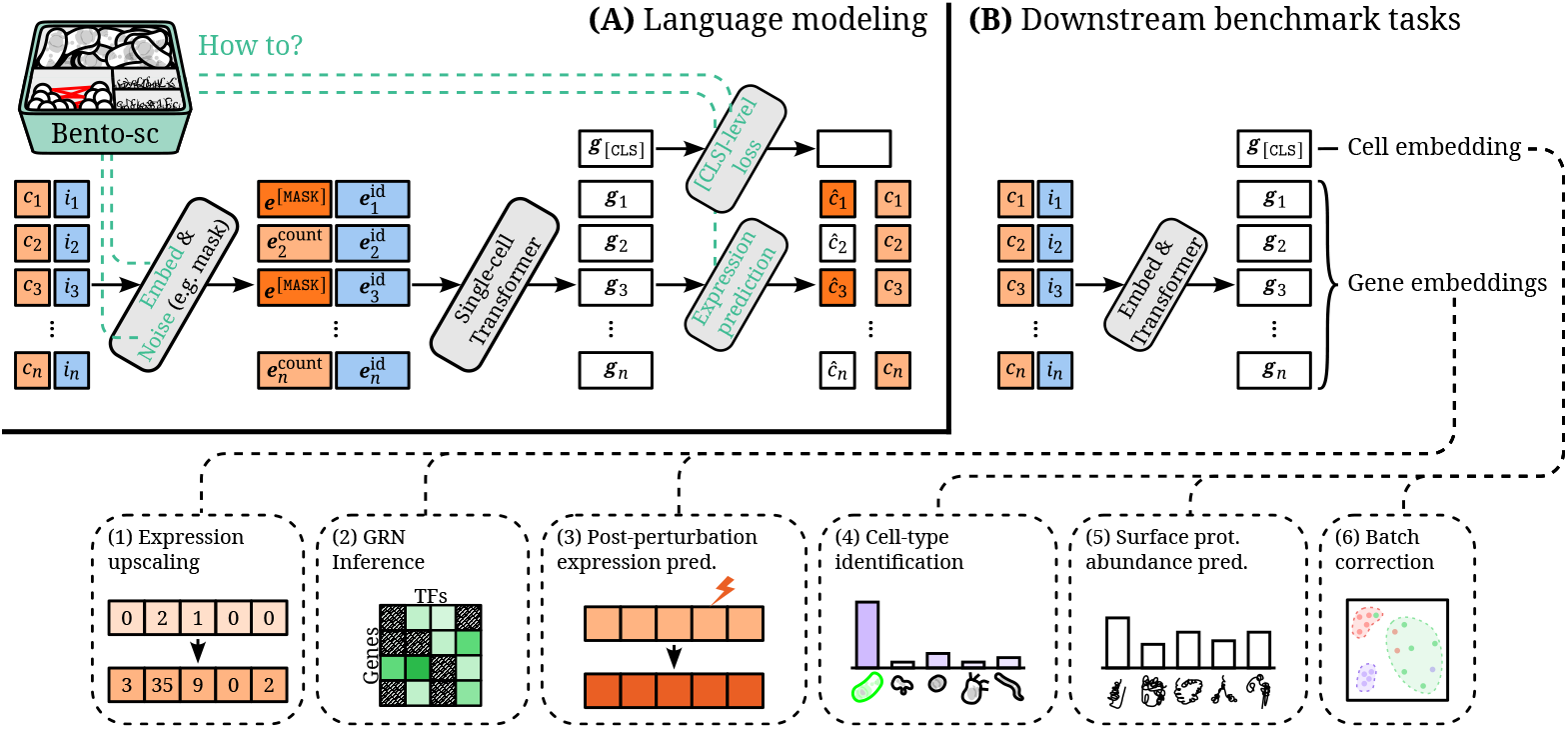
**(A)** Language model design tuning with *Bento-sc*. An input single-cell expression profile consists of non-zero counts *c*_*i*∈ {1,…,*n*}_ (orange), belonging to genes with indices *i*_*i*∈ {1,…,*n*}_ (blue). Both are embedded, and, after a stochastic masking step of count embeddings, are sent through a transformer. The transformer produces gene embeddings ***g***_*i*∈ _{_1,…,*n*}_ and a cell-level embedding ***g***_[CLS]_ . Both types of embeddings can be used in pre-training, through masked language modeling with an expression prediction head, and a `[CLS]`-level task, respectively. Bento-sc aims to tune four design choices of this pipeline, indicated in green, in order to obtain better performing single-cell language models. **(B)** Downstream benchmark tasks used for performance evaluation in this study. A summary of all tasks is given in Table 1.

The goal of *bento-sc* is to provide a framework to iterate upon this design, leading to an evaluation of scLM components (see Figure 1). Specifically, we tune variants of (1) the count embedding function, (2) the expression prediction reconstruction loss, (3) the noising (masking) function, and (4) the inclusion of loss functions on the cell-level (i.e. via ***g***_[CLS]_).

**Table 1.** Downstream evaluation tasks. Each task uses either gene (***g***_*i*∈ {1,…,*n*}_) or cell-level (***g***_[CLS]_) representations. Some tasks are fine-tuned on a training dataset, whereas others use the pre-trained representations as is (zero-shot /ZS).

| Task | Used Repr. | ZS | Performance Metric | Dataset(s) |
| --- | --- | --- | --- | --- |
| (1) Expression upscaling | Gene | ✓ | Pearson corr. | scTab<sup>[20]</sup> |
| (2) GRN inference | Gene | ✓ | $R^2$ | NeurIPS 2023 Perturbation<sup>[21]</sup> |
| (3) Post-perturbation expression pred. | Gene | ✗ | Pearson corr. | Perturb-seq<sup>[22]</sup> |
| (4) Cell-type identification | Cell | ✗ | Balanced accuracy | scTab<sup>[20]</sup> |
| (5) Surface prot. abundance pred. | Cell | ✗ | Pearson corr. | NeurIPS 2021 CITE-seq<sup>[23]</sup> |
| (6) Batch correction | Cell | ✓ | scIB score | (1) Zhang et al.<sup>[24]</sup>,<br>(2) Jorstad et al.<sup>[25]</sup>,<br>(3) Ivanova et al.<sup>[26]</sup> |

### A set of tasks for the evaluation of single-cell language models

In order to determine the capabilities of any scLM, it is insufficient to look at the pre-training by itself. This is because the performance on masked language modeling itself may not correlate with performance on various common single-cell prediction tasks. In this section, we lay out a diverse set of six tasks for evaluating scLMs: (1) expression upscaling, (2) GRN inference, (3) post-perturbation expression prediction, cell-type identification, (5) surface protein abundance prediction, and (6) batch correction (Table 1). This benchmarking line-up covers tasks that operate on cell-summary level tokens, as well as tasks that require predictions on the gene level. In addition, some tasks evaluate the learned representations of the model on novel tasks without fine-tuning (zero-shot), whereas other tasks involve fine-tuning with a separate training data set. A visual representation of every task is given in Figure 1. For details on implementation, data, and performance evaluation, see Methods.

All tasks use metrics with a maximum score of 1, where a higher score indicates a better model (see Table 1). To determine the overall quality of a scLM, we apply the model to each task, and compute the min-max scaled average performance across tasks. The min-max scaling is applied in such a way that the worst and best model on every task obtain a zero and one score, respectively. This operation makes it so that every task is weighted evenly, as the performance scores for one task may naturally occupy a larger numerical range than other tasks. We motivate the choice of “averaging performances across tasks” because this aligns with the general goal of foundation models: excelling in all biological tasks.

### Bento-sc: evaluating single-cell transformer configurations

Here we pre-train and evaluate a series of single-cell transformer language models. Our experiments can be roughly divided in four sections, each targeting a different component of single-cell transformer design. In every experiment, a series of transformer LMs are pre-trained that are identical to a “base” design except for one component. After each round of testing, the best-performing configuration is carried over to the next round. In other words, our study design follows that of greedy hyperparameter optimization at the scale of language models. We adopt this experimental setup due to the high computational cost of language model pre-training. Best performing models in each round are chosen via their performance on downstream tasks. In what follows, unless otherwise mentioned, performance on validation sets are shown.

#### Input count embeddings

In the first benchmarking step, we benchmark different choices for input count encoding functions 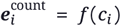 (Figure 2A). We start from a base language modeling design following the “general recipe” outlined above. Following scGPT^[12]^, the base design uses a count binning function. For *n* bins delimited by [*b*_1_, …, *b*_*n*+1_], a count *c* is assigned bin *i* according to: *c* ∈ [*b*_*i*_, *b*_*i*+1_)⇒ *i*. Bins are constructed in a way such that at least 0.2% of all counts occupy each bin^2^. Gene count embeddings are then obtained using a look up table similar to the gene identity embedding.

**Figure 2.**
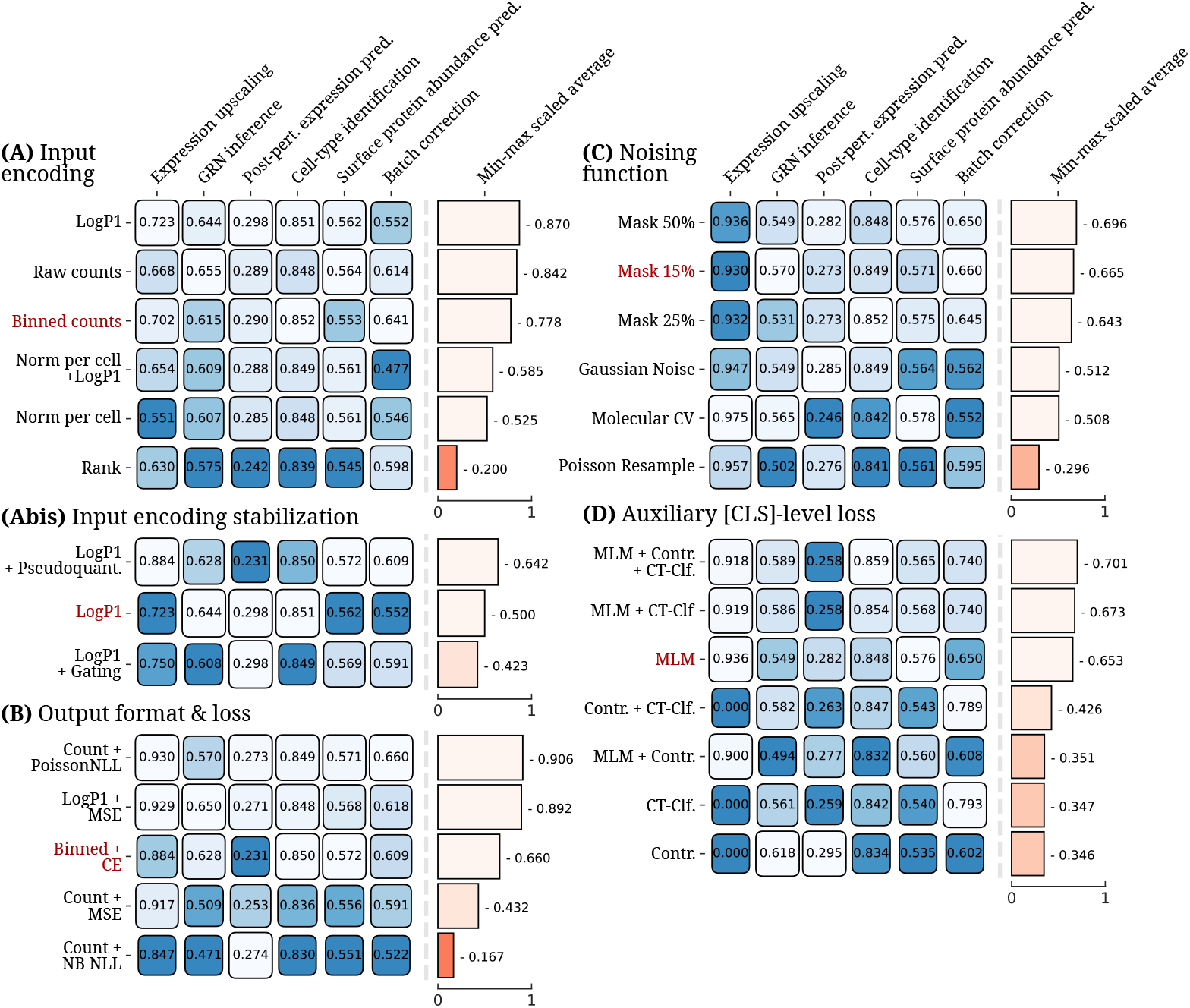
All main *bento-sc* validation results. **(A), (Abis), (B), (C)**, and **(D)** show results for testing different input encodings, input encoding stabilization tricks, count prediction formats and corresponding language modeling loss functions, MLM noising functions, and ***g***_[CLS]_ level loss functions, respectively. Lighter colors indicate better scores. For every experiment, the model indicated in red denotes the “base” default design, taken from the previous step. Min-max scaling for averaging task performance is always performed within experiment. Models are shown sorted by descending average performance.

As alternative gene input encoding strategies, we consider using (1) rank-based encoding (following Geneformer^[11]^), (2) raw counts, (3) normalization per cell (i.e. size factor normalization), (4) LogP1transformed counts, (5) normalization per cell + LogP1. The rank-based encoding involves ranking genes by expression level in descending fashion, assigning them their integer rank, and embedding the rank with a look-up table similarly as with binned counts. All other encoding strategies are well-known preprocessing operations in scRNA-seq^[27]^, and are here used in combination with a learnable linear projection layer ***W*** ∈ ℝ^1×*d*^. Note that in this experiment, while binned input counts are exchanged for alternative input strategies, we still predict binned counts at the output level.

After pre-training six single-cell language models, each using one of the six candidate input embedding strategies, they are evaluated on the six previously-mentioned downstream tasks (Figure 2A). It is observed that using LogP1-transformed or raw counts perform the best on average, with binned count input encodings not following far behind. Performing size factor normalization appears to hinder performance. Finally, rank-based input encoding is found to be worst-performing strategy.

To validate whether rank-based input encodings are unfairly disadvantaged in the context of trying to predict binned count levels at the output level, we perform an experiment using alternative output strategies. In the first, the binned output counts are exchanged for integer-valued ranks, equalizing the input and output encodings. In the second, gene identities are masked instead of count embeddings, and the model is pre-trained to predict which (masked) gene occupied the respective rank. This set-up closely resembles Geneformer’s design^[11]^. We found both alternative designs to be performing worse than using binned counts at the output level (Appendix D Figure 4).

As LogP1-transformed input counts are found to be the overall best-performing input encoding strategy, it is used in the remainder of experiments. In a second follow-up experiment, input encoding stabilization tricks are tested. These layers follow the input count embedding layer and are meant to stabilize the embeddings in case exceptionally large values are encountered. Here, two designs, which we will call “pseudoquantization” and “gating”, are adopted from Hao et al.^[14]^ and Clauwaert et al.^[28]^, respectively. Our experiments indicate that pseudoquantization helps overall results (Figure 2Abis). Hence, it will be adopted in the remainder of experiments.

#### Output format and loss

In a second step, output formats and loss functions are evaluated. These govern the supposed distribution we assume the counts to follow. As a base design, a model is trained to predict a count bin (constructed in the same way as the binned input strategy). This design involves an output linear layer ***W***_out_ that takes gene embeddings ***g***_*i*∈{1,…,*n*}_ ∈ ℝ^*d*^ and produces count bin probabilities ***ĉ***_*i*∈{1,…,*n*}_ = Softmax(***g***_*i*∈{1,…,*n*}_***W***_out_). These outputs are then optimized via the multi-class cross-entropy loss.

Alternative output formats and loss functions are to predict scalar counts, either raw or log-transformed. Raw count predictions are paired with the following loss functions: mean-squared error (MSE), Poisson negative log-likelihood (NLL), or negative binomial (NB) NLL, each assuming a different prior count distribution. Note that the two latter distributions cannot hold when zero counts are excluded. Instead, we model their zero-truncated counterparts (see Methods). LogP1 transformed count predictions are paired with the MSE loss. Figure 2B shows that the best performing scLMs result from those modeling either Poisson-distributed counts, or normally-distributed LogP1-transformed counts. As the former is the overall best performer by a small margin, it is adopted in the remainder of the experiments.

#### Noising functions

The third set of experiments consists of evaluating how inputs are corrupted for scLM optimization. The base design adopts masking at the same rate as Devlin et al.^[9]^. Here, we additionally test out two increased masking rates at 25% and 50%. In addition, input cell profiles can be corrupted in a different way than masking. Three additional strategies are evaluated: (1) adding Gaussian noise to the LogP1-transformed counts, (2) Using the counts as the mean *⇒* of a poisson distribution and resampling, and (3) performing molecular cross-validation^[29]^, which involves splitting each count into input and output fractions. These last methods have the additional advantage that they can be applied to all genes concurrently, and, hence, a pre-training signal is provided for all genes, rather than only the masked ones in MLM.

Our experiments show that MLM cannot be dethroned by alternative strategies (Figure 2C). While all three masking rates achieve similar results, a rate of 50% is found to the best performer, and hence is adopted in the remainder of the experiments.

#### Auxiliary [CLS]-level loss

As a last experiment, we call into question the added value of MLM as a whole. Alternative pre-training strategies entail contrastive learning and supervised pre-training, both operating on the cell summary embedding ***g***_[CLS]_ . In contrastive learning, a cell profile is corrupted (i.e. masked) twice independently to obtain two “views” of the same underlying cell. The model is then optimized to bring the cell embeddings ***g***_[CLS]_ of both closer to each other, while repelling all other cell embeddings^[30]^. In supervised pre-training, the transformer is trained to predict the cell-type from ***g***_[CLS]_, formulated as a multi-class classification task. As cell-type labels are available for large-scale datasets, such as the scTab dataset^[20]^, self-supervised modeling may not be necessary at all. As all these pre-training ways are non-mutually exclusive, we experiment with all their possible combinations.

Our experiments show that pre-training using either only contrastive or only cell-type classification results in generally worse performing models (Figure 2D). The top three models, all within a close small average performance margin, all use MLM. Among the top models, we observe that auxiliary [CLS]-level loss functions play a complementary function to MLM, amplifying performance. The best performing model uses all three discussed loss functions.

#### Final test performance and comparison with baselines

To conclude experiments, we compute test set results on all the models that arised as best performers within their experiment (Figure 3). This provides an unbiased view of the improvements obtained through each tuning experiment. In addition, the models are squared up against baseline methods for each task. Depending on the task, baselines can range from PCA, logistic regression, or unpre-trained transformers. Here, we only list the best baseline (selected on validation performance) per task (listed on Figure 3). For the full set of tested baselines, see Methods. For test performances, overall performance per model is additionally aggregated via computing the average rank across tasks.

**Figure 3.**
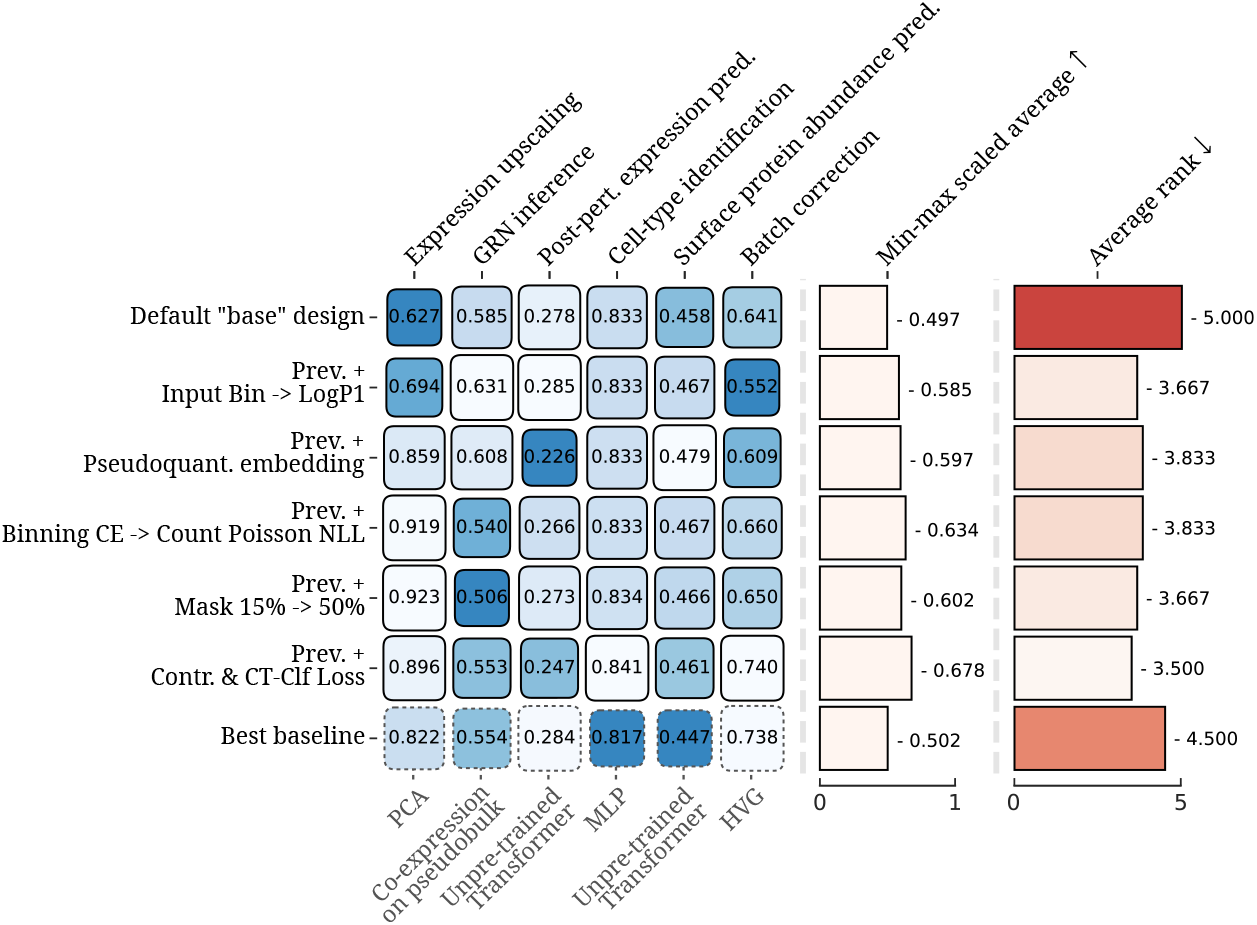
*Bento-sc* test results. Lighter colors indicate better scores. The best performers resulting from each tuning experiment are shown against best baselines per task. Models are shown sorted by chronological order of experiments, with baselines at the bottom. Here, model performances are shown aggregated by both minmax scaled averages, as well as average rank across tasks.

**Figure 4.**
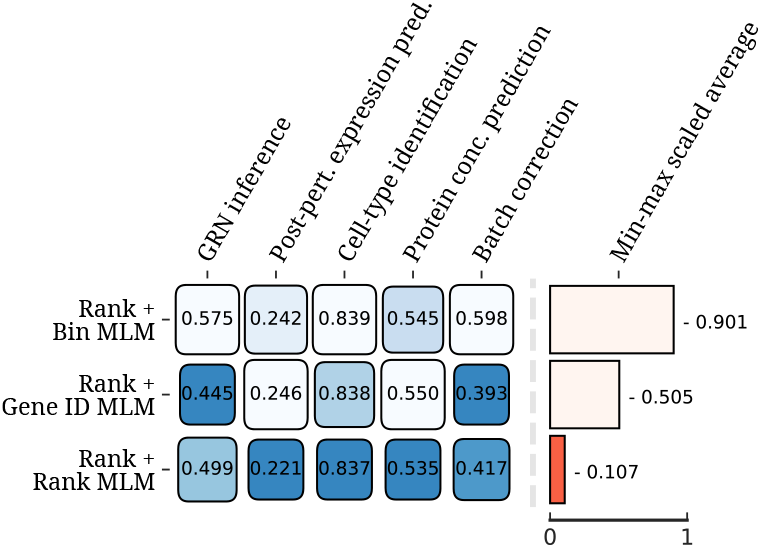
Evaluation of output loss functions for rank-based input encodings. Here, it is tested whether it is unfair to use the rank-based input encodings with binned output counts, as rank-based inputs may not allow to accurate predict count-level information. Instead of using binned output counts, here, the rankbased encodings is paired with either (1) trying to predict ranks at output level, and (2) masking Gene IDs and predicting them, both formulated as multi-class classification tasks, similar to MLM on binned counts. As the Gene ID MLM does not output counts, the Expression upscaling task is excluded from comparisons here.

**Figure 5.**
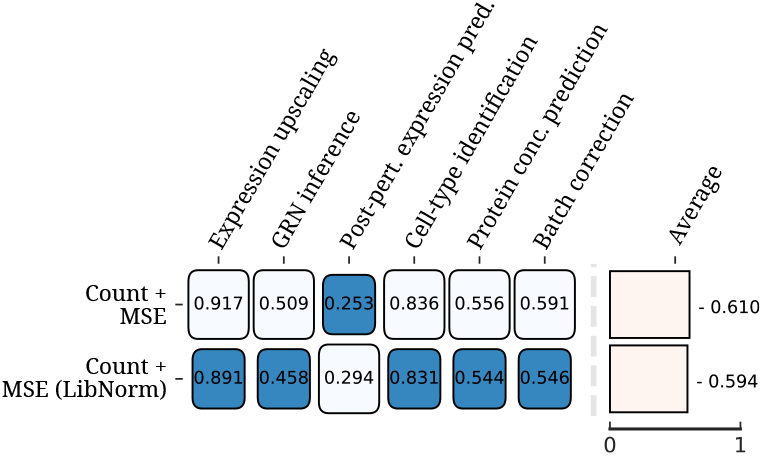
Evaluation of different instantiations of MSE Loss on raw counts (see Methods). Here, the simple average across tasks is taken, instead of the min-max scaled average. This is because the min-max scaled average computed across two models simply amounts to taking the average rank, losing all other quantitative information.

**Figure 6.**
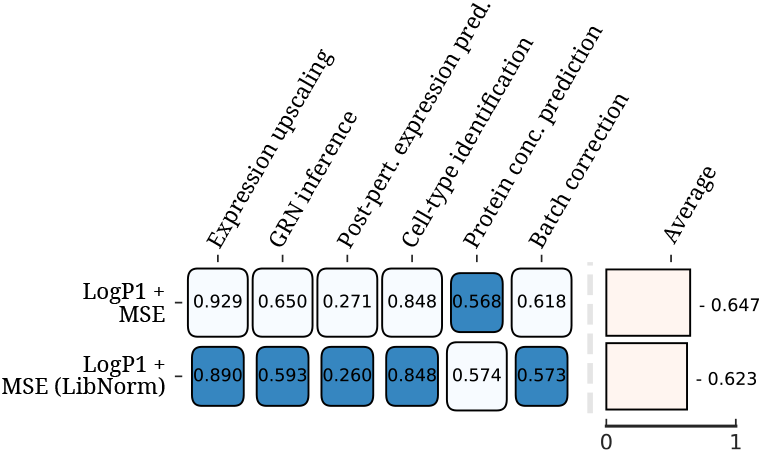
Evaluation of different instantiations of MSE Loss on LogP1 output counts (see Methods). Here, the simple average across tasks is taken, instead of the min-max scaled average. This is because the min-max scaled average computed across two models simply amounts to taking the average rank, losing all other quantitative information.

**Figure 7.**
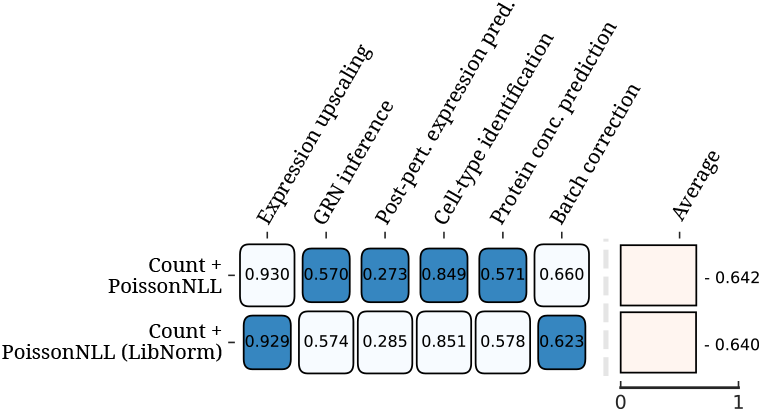
Evaluation of different instantiations of Poisson NLL loss on output counts (see Methods). Here, the simple average across tasks is taken, instead of the min-max scaled average. This is because the min-max scaled average computed across two models simply amounts to taking the average rank, losing all other quantitative information.

**Figure 8.**
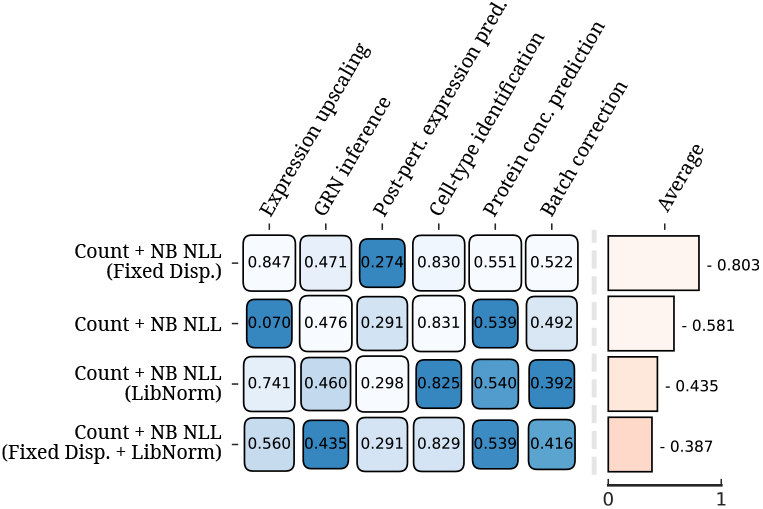
Evaluation of different instantiations of Negative Binomial NLL loss on output counts (see Methods).

**Figure 9.**
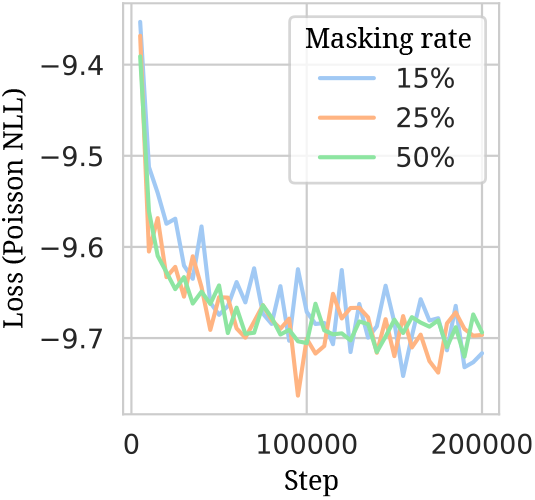
Pre-training validation loss curves for the three models using different masking rates.

From Figure 3, we can see that not every round of component tuning results in better test set performances. During tuning, performance on one task is often sacrificed for a better average performance across all tasks. It is also clear that no single language model is the best across all tasks. In general, however, the final model obtained through *bento-sc* is the best performer across tasks both in terms of min-max scaled averages, as well as average ranking. In addition, the final model never obtains the worst score on any task, signifying that it can successfully be employed towards any tested task. This objective is in line with the general aim of designing foundation models.

The best baselines for all tasks remain - on average - competitive with our line-up of scLMs. For two of the six tasks (cell-type identification and surface protein abundance prediction), baselines scored worse than all the language models shown in Figure 3. For two other tasks, Post-perturbation expression prediction and batch correction, baseline performance is virtually identical to the best performance we could obtain using scLMs. On average, baselines beat our default “base” language model design by a narrow margin. This advantage is not present for all other scLMs, which - on average - obtain superior performances. All models, however, occupy a small margin of performance differences. This is evident from the fact that the average rankings of all models are quite close to each other.

## Discussion

In this study, we systematically benchmarked design choices for constructing single-cell transformer language models, ultimately obtaining an optimized modeling recipe. Within every experiment, multiple design options were found to be roughly competitive. Hence, numerous ways to succesfully pre-train single-cell transformers exist. In the following paragraphs, we formulate some best practices.

The first two experiments concern the numerical presentation of counts to the model, at the input and output level, respectively. In general, minimally processing counts performs best, both at input and output level. Both rank-based and size factor normalized inputs perform substantially worse than simply using LogP1-transformed or raw counts. At the output, modeling raw counts under a Poisson distribution performs best, and is roughly on par with modeling LogP1-transformed counts as normally distributed. Hence, it is benefecial to employ prior knowledge on count distributions in loss function design. The exception to this guideline is the NB NLL loss, which performs the worst out of all tested reconstruction loss functions. We hypothesize that this is the case due to numerical instability issues w.r.t. its zero-truncated NLL (see Methods). It has to be noted that, to our knowledge, no prior works exist on incorporating Poisson NLL into single-cell transformer pre-training.

In the third experiment, we found that masking could not be dethroned by alternative noising functions. Masking remains effective up until high percentages, such as 50%. This may indicate that it is possible for a scLM to accurately reconstruct masked gene counts given only a fraction of the other expression values. Indeed, upon inspecting the pre-training loss curves for the different tested masking rates (Appendix D Figure 9), we find virtually no difference in reconstruction validation loss. Hence, further research needs to be dedicated to explore the limits of noising cell profiles for pre-training.

In the final experiment, it is observed that MLM pre-training generally outperforms contrastive and supervised pre-training. However, the two latter ways of pre-training were found to be complementary to MLM, as the best performer uses all three loss functions. The complementarity can be seen in the the performance for certain tasks. For example, the only models obtaining a batch correction score exceeding 0.70% are the ones that use cell-type identification supervised pre-training. This is expected, as separating cell-types into clusters is one component of the scIB score on which batch correction is assessed. On the other hand, without MLM as part of the pre-training procedure, the model cannot perform expression upscaling in a zero-shot fashion. A general conclusion is, hence: if one wants a single-cell foundation model to be applicable towards a wide variety of downstream tasks, the pre-training task should cover training signal that is relevant for all possible tasks.

Test set results reveal that unpre-trained baseline methods often remain competitive with pre-trained language models. Hence, pre-training does not offer the generational leap in prediction performances promised by foundation models. Further, not a single LM design outperforms baselines on all tasks. Consequently, scLMs cannot be realistically marketed as “one-stop shops” for single-cell analyses (yet).

There may be multiple reasons why pre-training does not enhance performance consistently. Here we put forward one hypothesis. In this study, pretraining was performed on the scTab dataset, which exclusively contains 10X-based data. Many of the downstream tasks involve input data gathered using non-10X-based sequencing protocols. For example, the surface protein abundance prediction and post-perturbation expression prediction tasks employ CITE-seq and Perturb-seq data, respectively. The transferability of models to other sequencing protocols has been previously studied in Fischer et al.^[20]^. There, a model trained on scTab obtains a macro F1-score of ±0.4 when evaluated on non-10X-based data, a drop from ±0.8. As each sequencing protocol comes with their own technical artefacts, this downstream data is essentially out-of-distribution (OOD). Hence, more research needs to be dedicated towards defining pre-training corpora that sufficiently cover the data manifold encountered in the many single-cell downstream tasks.

In conclusion, through iterative refinement, we identify successful (and failing) design choices for constructing scLMs. By carefully examining their performance against baselines, this study contributes to the growing body of research highlighting both the strengths and limitations of current scLMs. Via our PyPI-distributed Python package, *bento-sc* lays the groundwork for further experimentation with scLM design, and could be indispensable for future bench-marking approaches.

## Methods

### Pre-training experiments

#### Data

All pre-training experiments were performed on the scTab dataset^[4;20]^, derived from the CELLx-GENE^[31]^ census (version 2023-05-15). This dataset was chosen because: (1) it’s a long-term supported release, (2) original dataset splits are reproducible, and (3) the data is available in raw (unprocessed) count format. The 22.2 million cells in scTab are divided into training, validation and test fractions counting 15.2M, 3.5M, and 3.4M cells, respectively. To speed up model evaluation in the validation stage, an additional subset of 100 000 cells from the validation set was made. Unless otherwise noted, validation set results on scTab arise from this subset.

#### Pre-training model and set-up

The design of the encoder-only transformer blocks closely follows current state-of-the-art practices^[32]^. Namely, the blocks use pre-LayerNorms and GeLU gated linear units in the feedforward (FF) networks^[33;34]^. All models used 10 transformer encoder blocks, each with a hidden dimension of 512, 8 attention heads, and employing a dropout rate of 0.2 in the attention heads and between FF layers. The total number of trainable model parameters in the transformer encoder blocks is 51.9M. This figure does not count input gene count or identity embedding layers, as well as output heads such as gene expression prediction layers, or downstream task-specific heads.

All models were pre-trained in BFloat16 precision on scTab for 200 000 steps with the Adam optimizer^[35]^. A constant learning rate was applied over all training steps. The exact value of which is dependent on the scLM configuration (see below). Gradients were clipped to a norm of 1. All models were pre-trained on 2 GPUs using a per-GPU batch size of 192 (totaling 384). Assuming 1024 input gene tokens for every cell, all models were hence pre-trained with 393K tokens per batch. The total number of pre-training tokens, then, amounts to 78.6 billion (of which 15%, or 11.8B, provide training signal). Batches were constructed using bucket batching to minimize the masking token overhead (see Appendix A).

In what follows, all tested scLM configuration choices are described. A pre-training configuration was often subjected to an additional hyperparameter (such as, for example, individually tuned learning rate, depending on the loss). To tune such hyperparameters, a similar protocol was always used: a series of scLMs using different values for the hyperparameter were pre-trained for 2 500 steps. The best value was then chosen based on the best validation loss after 2 500 steps.

#### Input count embeddings

In total, six input count embeddings were tested. A count embedding method is defined as any approach *f* (·) that produces a vector-valued count embedding ***e***^count^ from a raw count *c*, including possible preprocessing steps.

The base design uses count binning, paired with embedding them using a look up table. For *n* bins delimited by [*b*_1_, …, *b*_*n*+1_], a count *c* is assigned bin *i* according to: *c* ∈ [*b*_*i*_, *b* _*i*+1_) ⇒ *i*. The input count embedding is then produced via ***e***^count^ = ***E*** [*i*, :], with ***E*** ∈ ℝ^*n*×*d*^ denoting a look-up embedding matrix with learnable weights. Bins are constructed in a way such that at least 0.2% of all counts occupy each bin. Many bins in the lower count ranges automatically cover more than 0.2% by including only a single count integer. Consequently, in total, *n* = 29 bins are used.

The rank-based input encoding follows a similar strategy to binning. Instead of *i* being designated by bin intervals, it is determined by the rank of the count within that cell, in descending order. i.e. the highest count in a cell profile is assigned 1, and so on. If multiple genes in a cell have identical counts, a random rank was assigned within the tie range.

All other input encoding strategies are of the form ***e***^count^ = *f* (*c*) = *p* (*c*) ***W***_in_, with *p* (·) a preprocessing function and ***W***_in_ ∈ ℝ^1×*d*^ a linear projection layer with learnable weights. For the raw count and LogP1 input encodings, *p* (·) takes the form *p*(*c*) = *λc, p*(*c*) = ln(*λc* + 1), respectively. A fixed normalization factor *λ* was introduced to ensure numerical stability, and tuned among possible values: {0.01, 0.05, 0.1, 0.5, 1, 5, 10} . For both models, a factor *λ* = 0.1 was selected. For the models using normalization per cell (with a subsequent LogP1 or not), *p* (·) takes the form 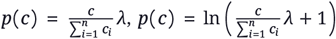, respectively. Again, the factor *λ* was tuned among possible values: {1e1, 1e2, 1e3, 1e4, 1e5, 1e6 }. For both models, a factor of *λ* = 1e3 was selected.

A single pre-training learning rate was used for all different input count embedding models. Following the protocol described above, the learning rate was tuned among possible values: {7e-5, 1e-4, **3e-4**, 5e-4}, with the best highlighted in bold.

Two input encoding stabilization “tricks” were also tested. These tricks manipulate the embedded count vector ***e***^count^ to make them robust to outlier counts. One trick, here denoted by “pseudoquantization”, was loosely adapted from Hao et al.^[14]^. Pseudoquantization processes a count embedding according to: Softmax (***e***^count^ ***W***_pq_), with ***W***_pq_ ∈ ℝ^*d*×*d*^ a learnable weight matrix. In other words, pseudoquantization uses the count embedding to create a weighted sum over rows in a learned matrix ***W***_pq_. Another tested trick, here denoted by “gating”, was adapted from Clauwaert et al.^[28]^. Gating processes a count embedding according to: tanh (***e***^count^)⊙ ***w***_g_, with ***w***_g_ ∈ ℝ^*d*^ a learnable weight vector.

#### Output format and loss

In total, five output format/loss combinations were tested. An output count format is defined by how it projects contextualized gene embeddings ***g***_*i*∈ {1,…,*n*}_ ∈ ℝ^*d*^ to expression predictions. It is then paired with possible suitable loss functions, based on output count format properties. In all cases, the gene reconstruction loss functions were computed as the mean loss over all masked tokens in the pre-training mini-batch.

As a default base design, binned output counts were used. The binning operation is identical to the one used for creating input count embeddings (i.e. using 29 different bins). Expression predictions were, hence, obtained as probability distributions over 29 bins. More formally, count predictions ***ĉ***_*i*∈{1,…,*n*}_ ∈ ℝ^29^ are produced according to: ***ĉ***_*i*∈{1,…,*n*}_ = Softmax(***g***_*i*∈{1,…,*n*}_***W***_out_), with ***W***_out_ ∈ ℝ^*d*×29^. For genes with masked input count embeddings, count bin predictions were optimized using the multi-class cross-entropy w.r.t. its true bin.

Alternative strategies predict raw counts, i.e. *ĉ*_*i*∈{1,…,*n*}_ ∈ [1, ∞), using *ĉ*_*i*∈{1,…,*n*}_ = exp (***g***_*i*∈ {1,…,*n*}_ ***w***_out_)+ 1, with ***w***_out_ ∈ ℝ^*d*^. The exponent and plus one ensure that predicted counts are left-bounded by 1. These count predictions were optimized to minimize the reconstruction error under three different count noise models: (1) normal (MSE loss), Poisson (Poisson NLL loss), and Negative Binomial (NB NLL loss). As, in this study, counts are left-bounded by 1, modified (i.e. zero-truncated) Poisson and Negative Binomial distributions are used. In both cases, *ĉ*_*i*∈ {1,…,*n*}_ ∈ [1, ∞) parameterizes the gene-wise mean of the estimated distribution. In the case of the NB distribution, an overdispersion parameter *θ*_*i*∈ {1,…,*n*}_ ∈ ℝ_>0_ is additionally estimated. We tested two ways to estimate *θ*_*i*∈{1,…,*n*}_: (1) as an extra output from ***g***_*i*∈ {1,…,*n*}_, similarly obtained via a learned projection (i.e. cell-specific dispersion), or (2) learning a fixed dispersion parameter vector 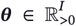, with *I* the total number of genes in scTab (19931). We found the fixed dispersion parameterization to perform better (Appendix D Figure 8). The full derivation of the zero-truncated Poisson and NB distributions and their negative-log likelihoods is given in Appendix B.

A final output format strategy is to predict LogP1-transformed counts, similarly obtained with a learned linear projection, and bounded to positive real numbers with an exponential function. The MSE loss was used to minimize the reconstruction error between predictions and LogP1-transformed ground truth counts.

For all strategies aside from the binned counts, an alternative realization of obtaining count predictions was tested. With this approach, the model predicted relative gene expression abundances (i.e. all count predictions were normalized to sum to one via a softmax operation). Count predictions were then obtained by multiplying the predicted relative abundances with the ground truth library size (correcting for the fact that counts are at a minimum 1). More formally, count predictions were obtained via

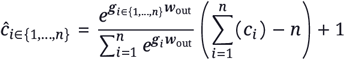

In all cases, this library-size normalized version of obtaining count predictions performed worse (Appendix D Figure 5,6,7,8).

All tested pre-training loss functions were subjected to learning rate tuning as described before. Table 2 shows learning rate tuning results.

**Table 2.** Learning rate tuning for pre-training different scLMs using different loss functions. Selected best learning rate are indicated in bold.

| Model | Learning rate |
| --- | --- |
| Count + MSE | {1e-5, 3e-5, 5e-5, 7e-5, 1e-4, **3e-4**, 5e-4, 7e-4, 1e-3} |
| Count + MSE (LibNorm) | {1e-5, 3e-5, 5e-5, 7e-5, **1e-4**, 3e-4, 5e-4, 7e-4, 1e-3} |
| Count + Poisson NLL | {1e-5, 3e-5, 5e-5, 7e-5, 1e-4, **3e-4**, 5e-4, 7e-4, 1e-3} |
| Count + Poisson NLL (LibNorm) | {1e-5, 3e-5, 5e-5, 7e-5, 1e-4, **3e-4**, 5e-4, 7e-4, 1e-3} |
| Count + NB NLL | {**1e-5**, 3e-5, 5e-5, 7e-5, 1e-4, 3e-4, 5e-4, 7e-4, 1e-3} |
| Count + NB NLL (Fixed $\theta$) | {1e-5, 3e-5, 5e-5, 7e-5, **1e-4**, 3e-4, 5e-4, 7e-4, 1e-3} |
| Count + NB NLL (LibNorm) | {1e-5, 3e-5, **5e-5**, 7e-5, 1e-4, 3e-4, 5e-4, 7e-4, 1e-3} |
| Count + NB NLL (LibNorm & Fixed $\theta$) | {1e-5, 3e-5, **5e-5**, 7e-5, 1e-4, 3e-4, 5e-4, 7e-4, 1e-3} |
| LogP1 + MSE | {1e-5, 3e-5, 5e-5, 7e-5, 1e-4, 3e-4, **5e-4**, 7e-4, 1e-3} |
| LogP1 + MSE (LibNorm) | {1e-5, 3e-5, 5e-5, 7e-5, 1e-4, 3e-4, **5e-4**, 7e-4, 1e-3} |

#### Noising functions

Six pre-training noising functions were tested. All noising functions are used to generate non-trivial pre-training objectives. During fine-tuning, the noising functions are no longer applied. Three of which used masking as in Devlin et al.^[9]^. Formally, masking proceeds as follows. Let 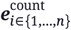 be the count embeddings for all *n* in a cell profile. A masking function produces noised count embeddings 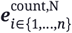, where:

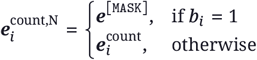

where *b*_*i*_ ∼ Bern (*p*) for all *i* ∈ {1, …, *n* }. Here, *p* denotes the masking probability. Following Devlin et al.^[9]^, the base design employs a masking probability of 15%. Alternative values of 25% and 50% were additionally tested. The reconstruction loss functions explained in the previous section are only applied for masked tokens.

Instead of masking a proportion of genes, alternative strategies were tested, which noise all genes in the cell profile at once. This has the advantage that the loss function can be applied to all gene tokens in all cells at any pre-training step. One such strategy involves noising the LogP1-transformed counts with Gaussian noise *ϵ* ∼ 𝒩 (0, 1) before they are embedded to higher-dimensional space. Another strategy involves noising the raw counts (before LogP1 transformation) by Poisson resampling. Poisson resampling uses the observed count *c*_*i*_ as the mean of a Poisson distribution, from which a sample is then taken to create 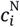, a noisy version of the input count: 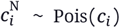, for all *i* ∈ {1, …, *n*}. If, after Poisson resampling, a count 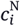 is turned to zero, the corresponding gene *i* was removed from the input and output expression profile.

The last tested noising strategy was molecular cross-validation^[29]^, which involves splitting each raw count (before LogP1 transformation) into input and output fractions. More formally, a noised input count 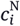 is a binomial subsampling of its observed count: 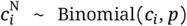, for all *i* ∈ {1, …, *n*}, with a probability *p*. Here, *p* = 0.10. The output count 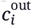, taken as ground truth to optimize reconstruction error against, is then taken as the difference: 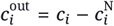. Genes for which either 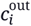 or 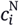 were turned to zero after molecular cross-validation, were removed from the input and output expression profiles. On average, expression profiles presented to the model using this strategy have a sequencing depth 10 times smaller than with other noising functions, effectively producing a data distribution shift. To offset this, the model using this noising function is pretrained on a subset of scTab containing all cell profiles with a sequencing depth of at least 25 000 total counts.

#### Auxiliary [CLS]-level loss

Two auxiliary loss functions on the cell-summary level were tested. All possible combinations of those two auxiliary losses with the MLM (gene expression reconstruction) loss were also tested. In those cases, the final loss was simply the sum of the individual loss terms. Note that for all models in this experiments, the masking noising function was always applied, even if the MLM loss was not included. As all models in this class concern modifications to loss functions, they were subjected to learning rate tuning as described before. Table 3 shows learning rate tuning results.

**Table 3.** Learning rate tuning for pre-training different scLMs using different auxiliary [CLS]-level loss functions. Selected best learning rate are indicated in bold.

| Model | Learning rate |
| --- | --- |
| Contr. | {1e-5, 3e-5, 5e-5, 7e-5, 1e-4, **3e-4**, 5e-4, 7e-4, 1e-3} |
| CT-Clf. | {1e-5, 3e-5, 5e-5, 7e-5, 1e-4, **3e-4**, 5e-4, 7e-4, 1e-3} |
| Contr. + CT-Clf. | {1e-5, 3e-5, 5e-5, 7e-5, 1e-4, **3e-4**, 5e-4, 7e-4, 1e-3} |
| MLM + Contr. | {1e-5, 3e-5, 5e-5, 7e-5, **1e-4**, 3e-4, 5e-4, 7e-4, 1e-3} |
| MLM + CT-Clf. | {1e-5, 3e-5, 5e-5, 7e-5, **1e-4**, **3e-4**, 5e-4, 7e-4, 1e-3} |
| MLM + Contr. + CT-Clf. | {1e-5, 3e-5, 5e-5, 7e-5, 1e-4, **3e-4**, 5e-4, 7e-4, 1e-3} |

**Table 4.** Validation performance of all baselines.

| Task | Method | Score |
| --- | --- | --- |
| Expression Upscaling | PCA | **0.860** |
| GRN Inference | Co-expression raw | 0.580 |
|  | Co-expression pseudobulk | **0.595** |
| Post-pert. expr. pred. | MLP-Mixer | 0.265 |
|  | Unpre-trained transformer | **0.295** |
| Cell-type identification | Logistic Regression | 0.764 |
|  | MLP | **0.833** |
|  | Unpre-trained transformer | 0.831 |
| Surface prot. abundance pred. | Logistic Regression | 0.462 |
|  | MLP | 0.540 |
|  | Unpre-trained transformer | **0.554** |
| Batch Correction | HVG 2000 | **0.738** |
|  | HVG 5000 | 0.735 |
|  | All genes | 0.730 |

One auxiliary [CLS]-level loss is supervised celltype identification, formulated as a multi-class classification task across the 164 classes included in scTab. Predicted cell-type probabilities are produced from cell-summary embeddings ***g***_[CLS]_ ∈ ℝ^*d*^ according to: ***ŷ***^ct^ = Softmax (***g***_[CLS]_***W***_ct_), with ***W***_ct_ ∈ ℝ^*d*×164^. The cross-entropy loss was used to optimize cell-type predictions.

An alternative self-supervised objective that operates on cell-summary embeddings ***g***_[CLS]_ is contrastive learning. In contrastive learning, a model first embeds two “views” of the same expression profile. The model is then optimized to bring those two embeddings close together in embedding space, while maximizing the dissimilarity with embeddings from other cells. Here, two views of the same cell are obtained by performing the noising function (i.e. masking at a rate of 50%) on a cell twice independently. This is performed for every cell in a mini-batch. Consequently, the batch size is doubled respective to its original size. In order to keep computational budgets constant, all models using contrastive learning are, hence, pretrained with a halved batch size w.r.t. the original cells included in the batch. More formally, a contrastive learning objective is constructed as follows: Within a mini-batch of *b* data points (i.e. cell profiles), each data point is noised twice to produce two “views”, forming a positive pair. All “views” are processed with a transformer to produce two cell summary embeddings 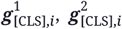 per cell *i*, with *i* ∈ {1, …, *b*}. The cell summary embeddings are then projected to a 128 dimensional contrastive embedding space via 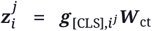, using a learnable weight matrix ***W***_ct_ ∈ ℝ^*d*×128^. We then apply the contrastive NT-Xent loss, as described in Chen et al.^[36]^, using a temperature of 0.1.

### Downstream tasks and baselines

In general, baseline method selection was informed by current best practices^[27]^. For the longevity’s sake of this benchmark, we favored the inclusion of simple, proven methods, such as a linear model for cell-type identification^[37]^, over more recent approaches.

#### Gene expression upscaling

For the gene expression upscaling task, no fine-tuning of pre-trained models was performed, i.e. it is a zero-shot task. We randomly select 25 000 deeply sequenced cells (>25 000 total counts) from the validation or test set in scTab (to produce validation or test set results, respectively). On these cells, we perform molecular crossvalidation performed in an identical way as for the pre-training set-up described above. To reiterate, an input count 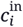 is obtained via binomial subsampling of the observed count: 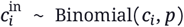, for all *i* ∈ {1, …, *n*}, with a probability *p*. Here, *p* = 0.10. The output count 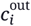, taken as ground truth to optimize reconstruction error against, is then taken as the difference: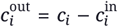. Genes for which either 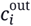 or 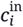 were turned to zero after molecular crossvalidation, were removed from the input and output expression profiles. Upscaling prediction quality is evaluated through the pearson correlation between predicted and ground truth expression profiles, computed across genes per cell and then averaged across cells. Not all models produce raw scalar counts as output of their pre-training task (i.e. the binned count and LogP1-transformed count output models). How these model predictions were converted to raw counts is discussed in Appendix C.

For this task, we included one baseline: a PCA model with 50 components, fitted on full 19331length gene expression profiles from the training set of scTab, which were library size normalized to 10 000 counts and LogP1-transformed. Gene expression predictions for evaluation were obtained on the same deeply sequenced and molecular crossvalidated scTab subset. Predictions were obtained from the PCA model by first applying the PCA transformation to principal component space, and then performed the inverse transformation. More formally, first: 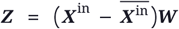, with input gene expression matrix ***X***^in^ ∈ ℝ^25000×19331^ and principal component matrix ***W*** ∈ ℝ^19331×50^, and then, to reconstruct: 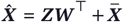 . The PCA baseline reconstructs expression levels of all genes, whilst the tested scLMs only reconstruct genes that are non-zero in both input and output profiles. To account for this fact, the pearson correlations between PCA predictions 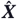 and molecular cross-validated output profiles ***X***^out^ were similarly only computed for the same subset of non-zero (input and output) genes.

#### Gene regulatory network (GRN) inference

GRNs were inferred from the pre-trained models without fine-tuning, i.e. in zero-shot fashion. They were obtained through the following procedure: First, an aggregated gene embedding 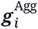 was computed for every gene *i* by averaging gene embeddings of that gene across a number of cells (here, 2500). A matrix ***S*** ∈ ℝ^*N*×*N*^, with *N* = 19331 the total number of genes, is then constructed using cosine similarities, i.e.: 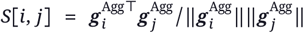 . This genegene correlation matrix was refined to a gene-TF matrix by using SCENIC’s cisTarget motif databases^[38]^. The goal is to filter the gene-TF matrix so that only regulatory targets of every TF are of non-zero similarity. After filtering, for every gene, the top-5 highest absolute similarity TFs are selected as regulators. A more detailed explanation GRN construction is given in Appendix C.

As baseline GRNs, a similar protocol as above was used. Instead of using output gene embeddings from a scLM, the gene-gene similarity matrix ***S*** was instead constructed from co-expression values. We tested two approaches: either co-expression on single-cell expression levels, or co-expression on pseudobulked expression levels. Pseudobulking was performed by grouping scTab profiles from the same donor. More formally, let ***X*** denote a matrix of single-cell or pseudo-bulked expression profiles consisting of full 19331-length gene expression profiles. All rows in ***X*** were library size normalized to 10 000 counts and LogP1-transformed. The gene-gene similarity matrix ***S***, was then filled with values according to: *S*[*i, j*] =*r*(***X*** [:, *i*], ***X*** [:, *j*]), with *r* the pearson correlation function. All subsequent steps to derive a final GRN, then, follow the same procedure as outlined in the previous paragraph. We found the pseudobulk-derived GRN to be slightly more performant (Appendix D Table 4).

As ground truth gene regulatory networks are largely undefined, how to properly evaluate inferred GRNs is an active field of research^[39]^. Here, we evaluate GRN inference through a proxy task. On a separate (pseudobulked) perturbation data set^[21;40]^, a linear regression task is constructed where gene expression levels of one gene is predicted using the expression levels of its (inferred) TFs as input features. Hence, the inferred GRN serves as a feature selection method. The intuition is that, the better the inferred GRN, the better their inferred TFs can predict expression levels in an external dataset.

To evaluate inferred GRN for multiple cell types, four separate GRNs were created for the following cell-types: (1) NK cells, (2) T cells, (3) B cells, and (4) Myeloid cells. For each of these cell types, 2 500 validation cell profiles were selected from scTab, either from its validation or test set, dependending on the evaluation stage. Separate cell-type specific GRNs were created following the procedures outlined above. Similarly, the regression proxy-task is also stratified per cell-type in the perturbation dataset, and hence, performed for four times. The final performance measure for this task is computed as the average ℝ^2^ score of the TF-to-gene linear regression models. A more detailed explanation of the regression proxy task implementation is given in Appendix C.

#### Post-perturbation expression prediction

The goal of post-perturbation expression prediction is to test whether the model can accurately predict expression profiles of cells with genetic perturbations unseen during training. As genetic perturbations (e.g. knock down or knock out of genes) may elicit a complex transcriptional response, it constitutes a non-trivial prediction task. Further, having a predictive model for unseen perturbations is practically useful, as it is infeasible to profile all possible permutations of perturbations.

This is a task for which the pre-trained models were fine-tuned on an external dataset. Here, we employed the Replogle et al.^[22]^ Perturb-seq dataset, also used in scGPT^[12]^. The Replogle et al.^[22]^ dataset contains cells from the K562 leukemia cell line, of which single genes were perturbed with CRISPR interference. To process the dataset, features (genes) were first mapped to the genes in scTab (assigning zero counts to all scTab genes not present in the original). In addition, cells that correspond to perturbations of genes not present in the remaining feature set were filtered out. After processing, the dataset contained 284 924 cells and 19331 genes. Of these 284 924 cells, 10 691 were control (unperturbed) cells. The remaining 274 233 cells were perturbed in one of 1866 possible genes. 80%/10%/10% of perturbations were assigned as training, validation, and test perturbations, respectively. Among the control cells, similarly 80%/10%/10% of cells were randomly assigned to training, val, and test sets. During training, every cell belonging to a training perturbation was randomly paired to a control cell during mini-batching. To ensure reproducibility, validation and test cells were deterministically paired with validation and test control cells, respectively.

To train a model to predict post-perturbation expression, pairs of control-perturbed cells were used. Given the control cell as input, the model was trained to predict the perturbed cell expression profile. Practically, a fixed 5000-gene space was constructed consisting of: (1) all genes with perturbations in the data, and (2) highly-variables genes (selected by descending variability until a set of 5000 genes was obtained). As this is a fixed gene set, it was, hence, possible for the model to encounter zero counts during fine-tuning. During fine-tuning, a “perturbation” indicator representation ***e***^pert^ was added to gene embeddings, communicating whether that gene was perturbed in the output state. Hence, for this task, input gene embeddings were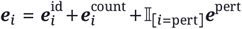. Fine-tuning details are given in Appendix C.

As baselines, two models were tested: (1) an MLP-Mixer^[41]^, and (2) a transformer using the exact same architecture as the final one, but not pre-trained on scTab. This last baseline allows to disentangle the benefit of pre-training vs architectural choices. We found the unpre-trained transformer to be more performant (Appendix D Table 4). Baseline details are given in Appendix C.

Following Cui et al.^[12]^, model performance was assessed through the perturbation-wise average Pearson_Δ_. This score measures the correlation between the predicted and true expression changes between post-perturbation and control cell states. Its calculation details are given in Appendix C.

#### Cell-type identification

Cell-type identification was formulated as a multi-class classification task. Here, we used the scTab dataset to fine-tune pretrained models towards this task. To prevent data leakage, the same cell training/validation/test splits were used as in pre-training. All reported validation set results arise from a 100 000 cell subset of the validation set.

All model weights were fine-tuned together with a linear classification output layer ***W***_ct_) ∈ ℝ^*d*×164^, which was trained from scratch to take the cell-summary embedding ***g***_[CLS]_ and to return cell-type logits ***ŷ***^ct^ = Softmax(***g***_[CLS]_ ***W***_ct_). The cross-entropy loss was used to optimize cell-type predictions. Model performance was evaluated through the balanced accuracy (equivalent to the macro-averaged accuracy). Fine-tuning details are given in Appendix C.

As baselines, three models were tested: (1) a logistic regression model, (2) an MLP, and (3) a transformer using the exact same architecture as the final one, but not pre-trained on scTab. This last baseline allows to disentangle the benefit of pre-training vs architectural choices. Among baselines, we found the MLP to be most performant (Appendix D Table 4). Baseline details are given in Appendix C.

#### Surface protein abundance prediction

Surface protein abundance prediction was formulated as a multi-output regression task. Paired single-cell expression and surface protein abundance measurements were obtained through CITE-seq data. Here, we used the NeurIPS 2021 CITE-seq^[23]^ dataset. This dataset comes with pre-configured train/val/test splits consisting of a test set with cells derived from a separate donor sequenced at a different site. To process the NeurIPS CITE-seq dataset, features (genes) were first mapped to the genes in scTab (assigning zero counts to all scTab genes not present in the original). After processing, the dataset contained 90 261 cells and 19 331 genes, with every cell additionally containing counts for 134 proteins. Of these 90 261 cells, 66 175 / 9 020 / 15 066 were training / validation / test cells, respectively.

All model weights were fine-tuned together with a linear classification output layer ***W***_prot_)∈ ℝ^*d*×134^, which was trained from scratch to take the cell-summary embedding ***g***_[CLS]_ and to return protein abundance predictions ***ŷ***^prot^ = ***g***_[CLS]_***W***_prot_. The model is trained to predict LogP1-transformed protein counts with the MSE loss. Model performance was evaluated through the macro-average pearson correlation. In other words, the pearson correlation between predicted and true LogP1-transformed protein counts was first computed for every protein target separately. Then, those pearson correlations were averaged to obtain a final score. Fine-tuning details are given in Appendix C.

As baselines, three models were tested: (1) a logistic regression model, (2) an MLP, and (3) a transformer using the exact same architecture as the final one, but not pre-trained on scTab. This last baseline allows to disentangle the benefit of pre-training vs architectural choices. Among baselines, we found the unpre-trained transformer to be most performant (Appendix D Table 4). Baseline details are given in Appendix C.

#### Batch correction

Batch correction was from the pre-trained models without fine-tuning, i.e. in zero-shot fashion. Here, we use three separate datasets derived from CELLxGENE, with no overlap with the scTab pre-training dataset. A first dataset consists of 195 632 circulating immune cells (either after COVID-19 infection, from vaccinated individuals, or healthy controls)^[26]^. Of the 195 632 cells in this dataset, 123 247 and 72 385 were profiled using 10x 5’ v1 and 10x 5’ v2 sequencing technologies, respectively. The second dataset is the human embryonic limb cell atlas^[24]^. Of the 125 955 cells in this dataset, 101 257 and 24 698 were profiled using 10x 3’ v2 and 10x 5’ v1 sequencing technologies, respectively. The third dataset is the human subset of a study profiling the middle temporal gyrus among great apes^[25]^. Of the 156 285 in this dataset, 141 782 and 14 503 were profiled using 10x 3’ v3 and Smart-seq v4 sequencing technologies, respectively. In each of these studies, the different sequencing chemistries were taken to be the batch effect. All of these datasets were processed by mapping their features (genes) to the genes in scTab (assigning zero counts to all scTab genes not present in the original). Note that this is the downstream task for which no distinction is made between validation and test stages.

For all of these datasets (separately), cell summary embeddings ***g***_[CLS]_ ∈ ℝ^*d*^ were obtained for every cell. Hence, per dataset, we obtained a matrix ***X*** ∈ ℝ^*n*×*d*^, with *n* and *d* the number of cells and scLM embedding dimension, respectively. This matrix was projected to a lower (50) dimensional space using PCA, yielding ***X***^PCA^ ∈ ℝ^*n*×50^ On this matrix, batch correction was performed via BBKNN^[42]^, a fast and lightweight batch correction algorithm. BBKNN produces a batch-corrected kNN graph of cells. Finally, the scIB score^[43]^ was then used to assess the batch correction performance. The scIB score balances batch correction quality with the conservation of biological information (measured through the tightness of cell-type clusters). The scIB score was calculated for all three datasets, and then averaged to produce the batch correction scores shown in this study. Details on the calculation of the scIB score is given in Appendix C.

As baselines, PCA was directly performed on (preprocessed) counts, instead of producing embeddings via a pre-trained model + PCA. In other words, the matrix ***X*** ∈ ℝ^*n*×*d*^ was now a preprocessed count matrix, instead of being filled with scLM cell embeddings. Counts were preprocessed via library size normalization to 10 000 counts and subsequent LogP1 transformation. Different values of *d* were tested: *d* ∈ {19331 (all), 5000, 2000 }. For *d* = 5000 or *d* = 2000, genes were selected based on their variability (i.e. selecting the *g*-most-variable genes in their respective datasets). All subsequent steps (i.e. PCA, BBKNN, and scIB calculation) were then performed identically as previously described. Among baselines, we found the version using the top 2000 highly variable genes (HVGs) to be most performant (Appendix D Table 4).

## Code and data availability

All code supporting this study is distributed on PyPI and is packaged under: https://github.com/gdewael/bento-sc. This study exclusively uses previously-profiled and openly accessible scRNA-seq data. All instructions to download data from their respective repositories and reproduce our experiments are given at https://github.com/gdewael/bento-sc.

## Acknowledgements

This work was supported by Research Foundation - Flanders (FWO) [PhD Fellowship fundamental research grant 1153024N to GDW]. WW also received funding from the Flemish Government under the “Onderzoeksprogramma Artificiële Intelligentie (AI) Vlaanderen” Programme The resources and services used in this work were provided by the VSC (Flemish Supercomputer Center), funded by the Research Foundation - Flanders (FWO) and the Flemish Government.

## Author contributions

All authors contributed in conceptualization of the study. GDW developed the software and performed the analyses. GDW wrote the original draft. GM and WW supervised the project. All authors contributed to discussions, interpreted the results, and revised the manuscript.

## A. On the exclusion of zero counts and FlashAttention for variable length data

Excluding all zero counts is a popular choice in single-cell language models^[10]^. On the surface, the reason for this is simple: at worst, self-attention’s memory and time complexity scales quadratically with input sequence length (as with naive self-attention computation). Even at best, the scaling remains linear (as with FlashAttention)^[19]^. As the vast majority of genes do not have observed counts in any single-cell expression profile, systematically excluding those from the input sequence allows practitioners to train bigger models using fewer resources. There is a second, more nuanced argument to make. In natural language, if we present a sentence to a transformer model, we only include input tokens for (sub)words actually occurring in that sentence. The act of not including all other words in the vocabulary communicates their absence in an implicit way. In a similar vein, if a gene input is not included to the transformer model, it may implicitly learn that this gene had no observed counts in that cell. This argument frames the choice as a biologically-motivated one, rather than one driven by pragmatism. The sensibility of excluding zero-counts, hence, relies on whether there is some innate biological signal in a zero-count that is not already adequately represented through the counts that *have* been observed.

In this work, we cap input sequence lengths at a maximum of 1024. This upper limit is useful because it means there is a predictable memory consumption during training—i.e. training will not suddenly halt due to an out-of-memory exception because an exceptionally deeply sequenced cell is encountered. All cell profiles with more than 1024 observed genes have some of their genes eliminated from the input space. We can minimize loss of information by eliminating those gene inputs with lowest counts first. To substantiate the sequence length cap of 1024, we present some figures: in our pre-training training data set, ∼71.25% of cells have more than 1024 observed genes. By putting a maximum of 1024 observed input gene tokens, we are throwing away ∼55.24% of total gene input tokens. Those thrown-out gene input tokens, however, represent only ∼25.91% of total counts in the data. If we assume that most biological meaning is present in the highly-expressed genes, enforcing a maximum input number of 1024 genes per cell presents a reasonable trade-off for computational efficiency with biological information retention.

Excluding zero counts turns our data into variable input length, as the number of observed genes will vary depending on the cell profile. By default, PyTorch’s implementation of FlashAttention does not support variable sequence lengths. The reason for this is that variable sequence lengths require padding tokens up to the maximum length if they are batched in a mini batch. To, then, make sure these padding tokens do not participate in computations, a selfattention mask is applied that prevents attention from padding tokens to “real” gene input tokens. It is this mask that is incompatible with current FlashAttention implementations. The FlashAttention repository provides an alternative “variable length” implementation of its algorithm, but its computational efficiency is lacking compared to “vanilla” FlashAttention^[44]^.

Here, we enable the usage of FlashAttention by performing the inverse of padding to the maximum sequence length in a batch: instead, we cut inputs to the minimum input length in a batch. We minimize loss of input data tokens by performing bucket batching (i.e. batching cells with similar number of input genes together). This procedure is implemented in a stand-alone python package https://github.com/gdewael/cut2min-bucket, and is discussed in more detail in a complementary blog post here: https://gdewael.github.io/blog/flashattnvarlen/.

## B. Zero-truncated count distributions and their negative loglikelihoods

### The zero-truncated Poisson distribution

The Poisson distribution is given by:

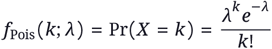

characterizing the probability of observing a (true) count *k*, given an (estimated) mean *λ*. Its zero-truncated version is, then:

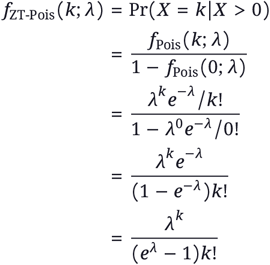

The negative log-likelihood of the zero-truncated Poisson distribution is, hence:

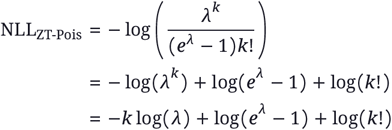

When estimating *λ* for an observed count *k*, the last term (log (*k*!)) can be ignored. Further, for large estimated numbers of *λ*, log(*e*^*λ*^ −1) becomes numerically unstable. As lim_*λ*→inf_ (log(*e*^*λ*^ − 1)) = *λ*, we can substitute the second loss term log(*e*^*λ*^ − 1) for *λ* when, say *λ* > 10.

### The zero-truncated Negative Binomial distribution

The Negative Binomial distribution is given by:

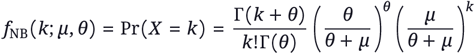

characterizing the probability of observing a (true) count *k*, given an (estimated) mean *⇒* and overdispersion *μ*. Its zero-truncated version is, then:

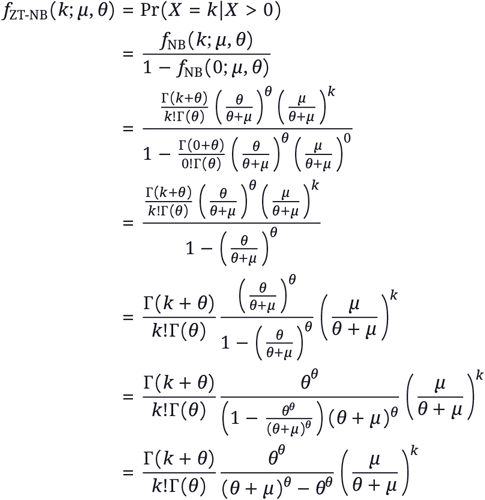

The negative log-likelihood of the zero-truncated Negative Binomial distribution is, hence:

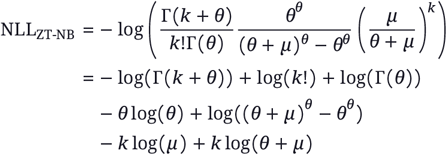

When estimating *μ* and *θ* for an observed count *k*, log(*k*!) can be ignored. Further, if *μ* and *θ* are both large, the term log ((*θ* + *μ*)^*θ*^ – *θ*)^*θ*^ becomes numerically unstable. In this case, we can show that:

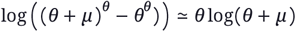

Hence, the one term can be interchanged for the other for large values of *μ* and *θ*. Here, we use the numerically stabilized term when either *μ* + *θ* > 15, or (*μ* – 1) (*θ* − 1)> 15.

## C. Implementation details of downstream tasks

### Estimating scalar counts with pre-trained models for gene expression upscaling

To compute pearson correlation between ground truth counts and predicted counts, the model needs to produce scalar-valued count predictions. All models in the first experiment use binned count outputs, i.e. predictions are obtained via ***ĉ***_*i*∈ {1,…,*n*}_ = Softmax (***g***_*i*∈ {1,…,*n*}_ ***W***_out_), with ***ĉ***_*i*∈ {1,…,*n* }_∈ ℝ^29^. In other words, these models predict a probability over count bins. To convert to scalar count predictions, the weighted sum over bins is taken: 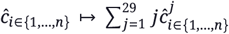 with *j* denoting a bin index.

To account for the fact that the predicted bin probabilities values might be uncalibrated (e.g. overconfident), the abovementioned procedure was performed using differently temperature-scaled softmax functions. For each model predicting binned counts, predicted bin probabilities were obtained via temperature-scaled softmax functions using a number of temperature values. For each temperature value, all predictions were converted to scalar using the aforementioned weighted sum function, and then used to produce a gene expression upscaling score. Hence, each model obtains multiple scores according to different temperature values ({0.01, 0.05, 0.1, 0.5, 1, 5, 10, 50, 100}). The best temperature for each model was always chosen on the validation set.

In experiment two, a model was tested that predicts LogP1 counts. Count predictions from that model were, hence, first corrected via *ĉ*_*i*∈ {1,…,*n*}_ ↦ exp(ĉ)−1, before being used to compute pear-_*i*∈{1,…,*n*}_ son correlation scores. All other tested model predict raw counts, which were produced directly from the pre-trained weights of the model.

### GRN construction

For each cell-type (NK cells, T cells, B cells, or Myeloid cells), 2500 cells were sampled from the scTab, either from its validation or test set, depending on the evaluation stage. Cells in scTab may have a more granular cell-type annotation than, for example, “B cell”. scTab includes many “sub-classes” (i.e. children in the cell ontology graph) of “B cell”, such as “mature B cell” and “immature B cell”^[45]^. Consequently, we sampled 2500 cells for each celltype, including all cells from cell-types which have the relevant cell type as parent class. From these cells, a gene-gene similarity matrix ***S*** ∈ ℝ^19331×19331^ was constructed, either using embeddings from a scLM, or using co-expression baselines, as described in Methods.

A first filtering on ***S*** filtering was to only consider genes that were observed more than 10 times across the 2500 cells. This was performed to limit spurious correlations. In addition, to eliminate self-connections in the GRN, all diagonal elements of ***S*** were put to 0. A further filtering step is performed by selecting for regulators of each gene. From SCENIC’s cisTarget resources^[38]^, a motif-gene matrix ***M*** ∈ ℝ^*m*×*g*^, with *m* and *g* the number of motifs and genes, respectively, was downloaded. Every entry of ***M*** represents the motif strength in the regulatory region of said gene (500bp upstream and 100bp downstream of its transcription start site). To select top motifs per gene, the matrix is first row-wise normalized according to the mean strength of every motif: 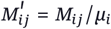, where 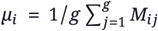. Then, the top 500 scoring motifs for every gene were selected as top hits. Motifs were mapped to TFs using a motif to TF database from SCENIC’s cisTarget resources. Consequently, for every gene, a list of TF regulators was obtained. The final GRN was then obtained by manipulating he gene-gene similarity matrix ***S*** so that, for every gene, only its TF regulators had a non-zero entry.

### GRN evaluation using regression proxy task

To evaluate GRN quality, an external (pseudobulked) perturbation dataset was used^[21;40]^. Prior to this study, the perturbation dataset was already batch corrected using scGen^[46]^ and normalized using pearson residuals^[47]^. In total, this dataset contained 2170 samples across 15215 genes, each sample belonging to one of 4 cell-types (NK, T, B, or Myeloid) and perturbed with one of 138 chemical compounds. Features (genes) in this dataset were first filtered for overlap with scTab. Then, 4 subsets of the dataset were created, selecting only cells from either of the tested cell-types (NK, T, B, or Myeloid cells). For each of these cell-types, in a previous step, a cell-type specific GRN was inferred (see above). In each cell-type specific subset, a regression task was constructed to predict gene expression levels from their regulators. Target to-predict genes were selected from the highly variable genes (HVGs) in the (cell-type subsetted) data. In the validation stage, we selected the top {1, 3, …, 199} HVGs as target genes. In the test stage, we selected the top {2, 4, …, 200} HVGs as target genes. For each of these genes, a linear regression model was trained to predict their expression levels from their TFs. To evaluate how well the GRN prioritizes top hits, for each gene, the 5 TFs with the highest absolute similarity score in ***S*** were selected. Every model was trained and evaluated on train and test fractions, consisting of a perturbation-split. 80% of the perturbations in the data were assigned to train, and the other to testing fractions. After training, an ℝ^2^ score was computed on the test fraction to reflect how well that gene could be predicted from its top identified TFs. These ℝ^2^ scores were averaged across all the HVGs that were selected as targets, to obtain a general score of the quality of the GRN for that cell-type This procedure was repeated for all cell-types, obtaining four average ℝ^2^ scores. These scores were, then, again averaged to obtain a final GRN Inference score.

### Post-perturbation prediction fine-tuning details

For post-perturbation prediction, one control-perturbed pair of cells constituted one training sample. The control cell was used as input, and the perturbed state was used as output. For both, a fixed gene set of 5000 genes were used as input genes (see Methods). At the output level, a new gene expression head was trained. In other words, the pre-training output layer that goes from gene embeddings ***g***_*i*∈ {1,…,*n*}_ ∈ **R**^*d*^ to count predictions *ĉ*_*i*∈ {1,…,*n*}_, was removed. Instead, a new output layer was put into place and trained from scratch. This output layer performs a similar function, going from gene embeddings to ***g***_*i*∈ {1,…,5000}_ to post-perturbation expression level predictions *ĉ*_*i*∈ {1,…,5000}_ . For this task, the models were always fine-tuned to predict perturbed cell states *c*_*i*∈ {1,…,5000}_ which were library size normalized to 10 000 counts and LogP1-transformed. The mean squared error loss was employed between true and predicted perturbed cell states.

All models were fine-tuned in BFloat16 precision with the Adam optimizer^[35]^. Fine-tuning was performed over a maximum of 50 000 steps, with validation performance checked every 500 steps. Finetuning was halted early when the validation Pearson_Δ_ did not improve over 40 validation checks. A constant learning rate of 7e-5 was applied over all training steps. Gradients were clipped to a norm of 1. All models were fine-tuned with a batch size of 32, with gradients accumulated over 4 batches (constituting a total batch size of 128).

The model uses 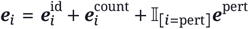 as input for a transformer model. ***e***^pert^ is initialized from ***e***^pert^ ∼ 𝒰(−1 /*γ*, 1 /*γ*)^*d*^, with *γ* a scaling factor. *γ* is tuned, together in a grid with the learning rate. Tuning was performed over a limited run of 5000 steps, with the best model selected on optimal validation Pearson_Δ_. Hyperparameters were tuned on the first (base) pre-trained scLM, and then used for all subsequent models. Tuned values for both are: *γ* ∈ {0.01, 0.1, **1**, 10} and lr ∈ {5e-5, **7e-5**, 1e-4, 3e-4, 5e-4}, with selected optimal values indicated in bold.

### Post-perturbation prediction baseline details

Here, a simple variant of the MLP-Mixer was used as baseline. Conceptually, this is one of the more simple models to still operate on genes as vector-valued tokens. Hence, it allowed to similarly incorporate a perturbation indicator token and enable generalization to unseen perturbations.

Let ***c*** ∈ ℝ^5000×1^ denote an input (control) cell profile, library size normalized to 10 000 counts and LogP1-transformed. Let *f*(·)= LayerNorm (Dropout(ReLU (·), 0.2)), a function applied after every linear layer in the MLP-Mixer. The whole model is, then, defined as

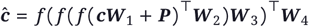

where 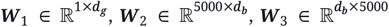, and 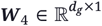 are learned linear projections in the MLP-Mixer. ***W*** _1_ embeds all genes to *d*_*g*_ dimensional space. ***P*** is a perturbation embedding matrix, filled as 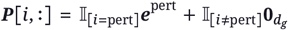, with 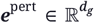 a learnable perturbation indicator vector (performing the same function as in the scLM). Parameters *d*_*g*_ and *d*_*b*_ are the number of hidden dimensions per gene and bottleneck dimension, respectively, controlling model complexity. ***W***_2_ and ***W***_3_ map to and from a *d*_*b*_ dimensional bottleneck space, respectively. Finally, ***W***_4_ maps back to (preprocessed) count space.

For MLP-Mixer, we considered the tuning of three hyperparameters: model size, batch size, and learning rate. As with the scLMs, ***e***^pert^ was initialized from 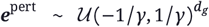, with *γ* = 1. These hyperparameters were tuned in a grid search using more limited runs of 5000 training steps. For the batch size *B*, the number of training steps *T* was scaled to retain equal compute budgets, according to: *T*_tune_ = 5000 /*B* · 128 and *T*_tune_ = 50000 /*B* · 128. The *B*-fold decrease of training steps with a *B*-fold increasing batch size is motivated by the perfect scaling regime studied by Shallue et al.^[48]^. The tested hyperparameters are: batch size *B* ∈ {**128**, 256, 512, 1024, 2048, 4096, 8192}, learning rate ∈ {1e-5, 3e-5, 5e-5, 7e-5, 1e-4, 3e-4, 5e-4, 7e-4, 1e-3, 3e-3, 5 e-3, 7e-3, **1e-2**}, and model sizes (*d*_*b*_, *d*_*g*_) ∈ {(32, 128), **(64**,**256)**, (128, 512), (256, 1024)}. Best values are indicated in bold. All other hyperparameter settings determining the training process were equal to the fine-tuning of the scLM models.

A second baseline consisted of an unpre-trained transformer. This variant used the exact same architectural set-up as the final model (i.e. LogP1 input encoding), but was initialized randomly at the start of perturbation modeling. In other words, it was not fine-tuned from a pre-trained scLM. All hyperparameters such as learning rate, batch size, and ***e***^pert^ initialization scaling factor *γ*, were all taken to be equal to those used in the scLM.

### Post-perturbation prediction performance evaluation

Model performance was assessed through the perturbation-wise average Pearson_Δ_. This score measures the correlation between the predicted and true expression changes between post-perturbation and control cell states. First, an average control cell profile is calculated over all *C* (val or test) control cells in the dataset:

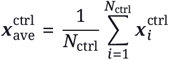

For every (val or test) perturbation *k* ∈ {1, …, *K*}, an average true 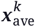 and predicted 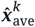 expression profile is similarly calculated:

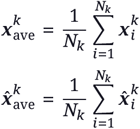

The post-perturbation expression change (Δ), is calculated over aggregated cell profiles: e.g. 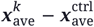. The Pearson_Δ_ is calculated as the average over perturbed genes *k*:

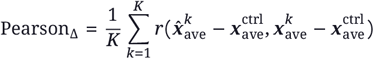

with *r* the pearson correlation function.

### Cell-type identification fine-tuning details

All models were fine-tuned in BFloat16 precision using the Adam optimizer^[35]^. Fine-tuning was performed with a batch size of 128 over a maximum of 500 000 steps, with validation performance checked every 5000 steps. Fine-tuning was halted early when the validation balanced accuracy did not improve over 40 validation checks. A constant learning rate of 3e-4 was applied over all training steps. Gradients were clipped to a norm of 1.

The fine-tuning learning rate was tuned on the first (base) pre-trained scLM, and then used for all subsequent (pre-trained) models. Tuning was performed over a limited run of 50 000 steps, with the best model selected on optimal validation balanced accuracy. Tested learning rates were {3e-5, 5e-5, 7e-5, 1e-4, **3e-4**, 5e-4, 7e-4 }, with the selected optimal value indicated in bold.

### Cell-type identification baseline details

For the logistic regression and MLP cell-type identification models, input counts were library size normalized to 10 000 counts and LogP1-transformed. The logistic regression model predicts cell-types via a single linear transformation ***ŷ***^ct^ = Softmax(***cW***_ct_), with ***W***_ct_ ∈ ℝ^*g*×164^, with *g* the number of input genes. The MLP uses multiple such linear transformations, with non-linearities inbetween. Let *f* (·) = ReLU(LayerNorm(Dropout(·, 0.2))), a function applied after every linear layer in the MLP. The whole MLP model is, then, defined as

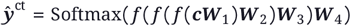

where ***W***_1_ ∈ ℝ^*g*×512^, ***W***_2_ ∈ ℝ^512×256^, ***W***_3_ ∈ ℝ^256×128^, and ***W***_4_ ∈ ℝ^128×164^ are learned linear projections in the MLP. The cross-entropy loss was used to optimize cell-type predictions.

For both the logistic regression and MLP models, we considered the tuning of three hyperparameters: number of input genes *g*, batch size, and learning rate. These hyperparameters were tuned in a grid search using more limited runs of 50000 training steps. For the batch size *B*, the number of training steps *T* was scaled to retain equal compute budgets, according to: *T*_tune_ = 50000/*B* · 128 and *T*_tune_ = 500000/ *B* · 128. The *B*-fold decrease of training steps with a *B*-fold increasing batch size is motivated by the perfect scaling regime studied by Shallue et al.^[48]^. The tested hyperparameters are: batch size *B* ∈ {128, 256, 512, ***1024***, 2048, 4096}, learning rate ∈ {1e-4, 3e-4, 5e-4, **7e-4**, 1e-3, 3e-3, 5e-3, *7e-3*, 1e-2, 3e-2, 5e-2, 7e-2, 1e-1}, and number of input genes *g* ∈ {***19331***, 5000, 2000}. Best values for the logistic regression and MLP models are indicated in bold and italic, respectively. For number of input genes *g* = 5000 or *g* = 2000, genes were selected based on their variability (i.e. selecting the *g*-most-variable genes). All other hyperparameter settings determining the training process were equal to the fine-tuning of the scLM models.

A third baseline consisted of an unpre-trained transformer. This variant used the exact same architectural set-up as the final model (i.e. LogP1 input encoding), but was initialized with random weights at the start of cell-type identification training. In other words, it was not fine-tuned from a pre-trained scLM. All hyperparameters such as learning rate and batch size were all taken to be equal to those used in the scLM.

### Protein concentration prediction fine-tuning details

All models were fine-tuned in BFloat16 precision using the Adam optimizer^[35]^. Fine-tuning was performed with a batch size of 128 over a maximum of 200 000 steps, with validation performance checked every 2000 steps. Fine-tuning was halted early when the validation macro pearson correlation did not improve over 40 validation checks. A constant learning rate of 1e-4 was applied over all training steps. Gradients were clipped to a norm of 1.

The fine-tuning learning rate was tuned on the first (base) pre-trained scLM, and then used for all subsequent (pre-trained) models. Tuning was performed over a limited run of 20 000 steps, with the best model selected on optimal validation macro pearson. Tested learning rates were {1e-5, 3e-5, 5e-5, 7e-5, **1e-4**, 3e-4, 5e-4, 7e-4, 1e-3}, with the selected optimal value indicated in bold.

### Surface protein abundance prediction baseline details

For the logistic regression and MLP baselines, an identical set-up was used as with cell-type identification. Input counts were library size normalized to 10 000 counts and LogP1-transformed. The only difference is that the output of the model is now 134 dimensional instead of 164, that there is no softmax operation, and that the MSE loss is computed. The MSE loss optimizes the model towards predicting LogP1-transformed protein marker counts. For logistic regression and MLP model details, see the section above discussing them within the context of cell-type identification.

For both the logistic regression and MLP models, the same three hyperparameters were tuned: number of input genes *g*, batch size, and learning rate. These hyperparameters were tuned in a grid search using more limited runs of 20000 training steps. For the batch size *B*, the number of training steps *T* was scaled to retain equal compute budgets, according to: *T*_tune_ = 20000/*B* · 128 and_tune_ *T*_tune_ = 200000 /*B* · 128. The *B*-fold decrease of training steps with a *B*-fold increasing batch size is motivated by the perfect scaling regime studied by Shallue et al.^[48]^. The tested hyperparameters are: batch size *B* ∈ {*128*, **256**, 512, 1024, 2048, 4096}, learning rate ∈ {1e-4, **3e-4**, 5e-4, 7e-4, 1e-3, 3e-3, 5e-3, *7e-3*, 1e-2, 3e-2, 5e-2, 7e-2, 1e-1}, and number of input genes *g* ∈ {***19331***, 5000, 2000} . Best values for the logistic regression and MLP models are indicated in bold and italic, respectively. For number of input genes *g* = 5000 or *g* = 2000, genes were selected based on their variability (i.e. selecting the *g*-most-variable genes in the NeuRIPS 2021 dataset). All other hyperparameter settings determining the training process were equal to the fine-tuning of the scLM models.

A third baseline consisted of an unpre-trained transformer. This variant used the exact same architectural set-up as the final model (i.e. LogP1 input encoding), but was initialized with random weights at the start of surface protein abundance prediction training. In other words, it was not fine-tuned from a pre-trained scLM. All hyperparameters such as learning rate and batch size were taken to be equal to those used in the scLM.

### scIB score calculation

The scIB score is a weighted sum of two components:

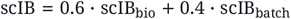

In this study, the bio-conservation score scIB_bio_ is calculated as the mean of four metrics: ARI, NMI, Isolated F1, and cLISI. The batch correction score scIB_batch_ is calculated as the mean of two metrics: Graph Connectivity and iLISI. These metrics include all the possible metrics under the scIB framework that are (1) possible to compute on batch-corrected kNN graphs, (2) are purely Python-based. For more details on all the individual metrics making up the scIB score, we refer the reader to Luecken et al.^[43]^.

## D. Figures and tables supporting the methods and results sections

1 While there is some work that adapts autoregressive transformers for single-cell data ^[18]^, the non-sequential nature of the genewise count data makes the masked language modeling (MLM) objective a more natural fit. In addition, it can be argued that autoregressive modeling is a special case of masked autoencoding, in which the masked tokens occur strictly at the end of the inputs.

2 In practice, equal frequency binning is impossible for scRNAseq data, as a bin containing only counts equal to one would cover ∼56.6% of all counts in scTab.

